# Discovery of pharmaceutically-targetable pathways and prediction of survivorship for pneumonia and sepsis patients from the view point of ensemble gene noise

**DOI:** 10.1101/2020.04.10.035717

**Authors:** Tristan de Jong, Victor Guryev, Yury M. Moshkin

**Affiliations:** European Research Institute for the Biology of Ageing, University of Groningen, University Medical Centre Groningen, Groningen, The Netherlands; Federal Research Centre, Institute of Cytology and Genetics, SB RAS, Novosibirsk, Russia; Institute of Molecular and Cellular Biology, SB RAS, Novosibirsk, Russia; Gene Learning Association, Geneva, Switzerland

## Abstract

Finding novel biomarkers for human pathologies and predicting clinical outcomes for patients is rather challenging. This stems from the heterogenous response of individuals to disease which is also reflected in the inter-individual variability of gene expression responses. This in turn obscures differential gene expression analysis (DGE). In the midst of the COVID-19 pandemic, we wondered whether an alternative to DGE approaches could be applied to dissect the molecular nature of a host-response to infection exemplified here by an analysis of H1N1 influenza, community/hospital acquired pneumonia (CAP) and sepsis. To this end, we turned to the analysis of ensemble gene noise. Ensemble gene noise, as we defined it here, represents a variance within an individual for a collection of genes encoding for either members of known biological pathways or subunits of annotated protein complexes. From the law of total variance, ensemble gene noise depends on the stoichiometry of the ensemble genes’ expression and on their average noise (variance). Thus, rather than focusing on specific genes, ensemble gene noise allows for the holistic identification and interpretation of gene expression disbalance on the level of gene networks and systems. Comparing H1N1, CAP and sepsis patients we spotted common disturbances in a number of pathways/protein complexes relevant to the sepsis pathology which lead to an increase in the ensemble gene noise. Among others, these include mitochondrial respiratory chain complex I and peroxisomes which could be readily targeted for adjuvant treatment by methylene blue and 4-phenylbutyrate respectively. Finally, we showed that ensemble gene noise could be successfully applied for the prediction of clinical outcome, namely mortality, of CAP and sepsis patients. Thus, we conclude that ensemble gene noise represents a promising approach for the investigation of molecular mechanisms of a pathology through a prism of alterations in coherent expression of gene circuits.

## Introduction

Both viral and bacterial pneumonia may lead to a life-threatening condition, namely sepsis. Most notable cases, in the public perception, include pandemic viral infections, such as the 2009 swine flu pandemic caused by H1N1 [1] and more recently, the 2019 coronavirus disease (COVID-19) caused by severe acute respiratory syndrome coronavirus 2 (SARS-CoV-2) [2]. Like with any other annual severe acute respiratory infections (SARI), these pandemics resulted in a significant raise in patients with sepsis at intensive care units[3, 4]. Sepsis is a complex reaction of the host (human) to a systemic infection (viral or bacterial) often resulting in septic shock or death [5-7]. A problem of sepsis treatment, the prediction of patients’ clinical outcomes and the risks of mortality relates to the highly heterogenous nature of sepsis [8]. Thus, despite recent progress in identification of molecular biomarkers for sepsis [8-15], treatment remains mainly non-curative and clinical outcomes are mostly inferred from clinical signs [5].

A canonical approach for the identification of disease biomarkers and their potential therapeutic targets relies on differential gene expression (DGE) analysis either on RNA or protein levels. This stems from a classical gene regulation Jacob-Monod model, which implies a specific gene expression response (up- or down-regulation) to a specific signal (see recent perspective on historical origins of the model in [16]. However, gene expression is a stochastic process and cellular responses to signals often trigger a cascade of changes in gene expression, making it difficult to discover specific targets and biomarkers for a disease.

The stochastic nature of gene expression implies a natural variation in RNA and protein copy numbers [17]. According to the fluctuation-response relationship [18, 19], an amount of gene expression response to a signal (fluctuation) is proportional to its variance (or squared biological coefficient of variation – *bcv*^*2*^) for log-scaled values of RNA copy number [20]. Consequently, statistical inference of differentially expressed genes will be biased towards genes with high variance (*bcv*^*2*^) (Figure S1). This leads to a set of intrinsic problems with DGE analysis. 1) genes with increased variability in expression will strongly respond to any cellular signal aimed at them. However, these genes may not necessarily be causative for a diseased state. Even under normal circumstances they exhibit large fluctuations and, thus, are loose-regulated. 2) In contrast, genes with a low variability will respond only modestly, but these genes are tight-regulated and any fluctuations in their expression might be causative for a diseased state.

Upon calling significantly changed genes, to make biological sense, these genes are mapped to known biological pathways, such as GO or KEGG [21, 22], or to subunits of protein complexes annotated by CORUM or other interaction databases [23]. Thus, a second statistical test is required, namely gene set enrichment analysis (GSEA). However, this is not without its own caveats. The major one is that GSEA depends on the statistical inference of DGE and DGE cut-offs [24, 25]. As a result, biological interpretations from DGE might be drastically affected by pitfalls arising from the fluctuation-response relationship, DGE thresholding and the choice of statistical approach for GSEA.

To circumvent this, we reasoned that 1) genes do not function in isolation, but rather act as ensembles representing biological pathways and/or subunits of protein complexes. 2) The normal function of a biological pathway or protein complex requires a regulated (balanced) expression of the whole gene ensemble. 3) Any alterations in the expression of a gene ensemble might be causative for a disease or predictive for clinical outcome. To infer the alterations in gene ensembles expression we turned to the estimation of their variances (ensemble gene noise) from whole blood gene expression profiles of individuals under normal and pathological conditions. From the total law of variance, ensemble gene noise (Var[*G*]) sums from the variance of ensemble genes’ means and (Var[E[*G*|*g*]]), and the expectation of ensemble genes variances (E[Var[*G*|*g*]]) (Figure S2). Thus, the ensemble gene noise estimates both: 1) changes in stoichiometries of genes encoding either a biological pathway or protein complex subunits and 2) changes in mean gene expression variability for genes in ensemble.

From the whole blood expression profiles of patients under intensive care treatment we estimated how ensemble gene noise corresponds to a pathological state, such as sepsis, community/hospital acquired pneumonia (CAP) or viral H1N1 pneumonia (H1N1). From this analysis we identified a number of pathways for which ensemble gene noise associated positively with an individual health/disease state treated as an ordinal variable (healthy < early H1N1 phase < late H1N1 phase and healthy < sepsis/CAP survived < sepsis/CAP deceased patients). Finally, we identified pathways and complexes where deregulation is associated with a poor prognosis and predicted the clinical outcome (survival/mortality) for CAP/sepsis patients based on ensemble gene noise with high accuracy. We concluded that the ensemble gene noise provides a powerful tool for the discovery of systemic disease biomarkers, pharmaceutically targetable pathways and the prediction of a disease clinical outcome.

## Results

### Mean and variance gene expression response to infection and sepsis

Sepsis is thought to trigger a plethora of heterogenous host responses to a systemic infection [5, 8]. We reasoned that this heterogeneity might be reflected in the inter-individual gene expression variability (standard deviation - σ or variance - σ^2^). Considering that a) RNA copy number is a mixed Poisson (*e.g.* negative binomial) random variable [26] and that b) log-transformed microarray hybridization signal intensities correlate with log-transformed RNA-seq copy numbers [27]. It is easy to show that the variance of log gene expression approximates the biological coefficient of variation (bcv^2^) [20]. From the first-order Taylor expansion for variance: 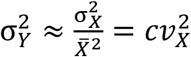, where *Y* = *log*(*X*) is the log gene expression. The mixed Poisson random variable, 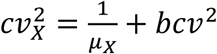, where *bcv*^*2*^, also known as the overdispersion parameter, is independent of mean gene expression (*μ*_X_). Thus, for *μ*_X_ ≫ 1 (for genes with a large mean RNA copy number), 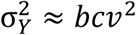. In other words, by estimating the inter-individual log gene expression variabilities from either microarray signal intensities or RNA-seq counts we can infer approximately the biological coefficients of variations for genes’ RNA copy numbers.

We estimated the dispersions for whole blood log gene expressions in CAP and sepsis patients (8826 genes), and H1N1 infected patients (7240 genes) from the two data sets GSE65682 and GSE21802 respectively (for a detailed description of cohorts see original studies and Methods) [8, 11, 28]. For CAP and sepsis patients we also accounted for age, including it as a random variable in the Generalized Additive Model for Location, Scale and Shape (GAMLSS) [29], see Methods. On average, the dispersions in log gene expressions in CAP, sepsis and H1N1 patients were significantly higher as compared to healthy individuals (Figure 1A). To that, for CAP patents’ dispersions in log gene expressions were significantly higher for deceased patients as compared to those survived. Likewise, for H1N1 patients, dispersions in log gene expressions further increased in the late phase of infection (Figure 1A). For sepsis patients, on average dispersions in log gene expressions were comparable between survived and deceased patients for all analysed genes (Figure 1A). However, for genes for which dispersions changed significantly between healthy individuals and sepsis patients (Bonferroni adjusted p ≤ 0.05), their dispersions on average were higher in the deceased patients as compared to the survived (p < 0.001). Together, these suggest that host response to infection increases the biological coefficients of variations in genes’ RNA copy numbers (as σ^2^ ≈ *bcv*^2^) and substantiates heterogeneity in the pathogenesis of sepsis [8] from the gene expression perspective.

**Figure 1.**
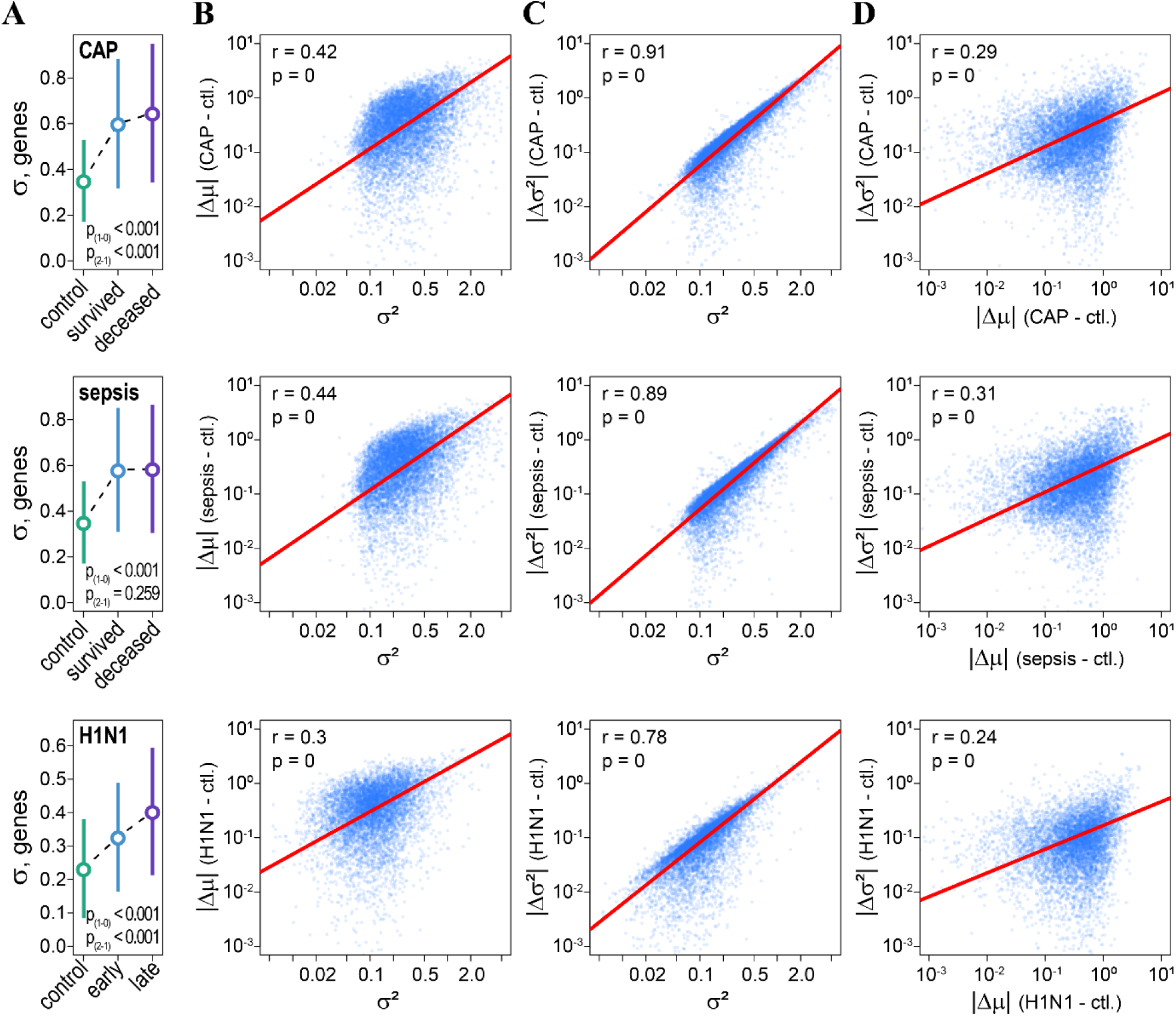
H1N1, CAP and sepsis coordinately affect mean gene expression and inter-individual gene expression variability. **A)** Inter-individual variability in whole blood gene expression (σ) increases in CAP (top), sepsis (mid) and H1N1 (bottom) patients as compared to healthy individuals. p_(1-0)_ – *p*-values of *t* tests comparing differences in inter-individual gene expression variability of healthy individuals (control) with survived (CAP, sepsis) and early H1N1 infected patients. p_(2-1)_ – *p*-values of *t* tests comparing differences of survived (CAP, sepsis) and early H1N1 infected patients with deceased (CAP, sepsis) and late H1N1 infected patients. Circles and whiskers indicate means and standard deviations respectively. **B)** Correlations between variances in whole blood gene expression (σ^2^) and absolute changes in mean gene expression (|Δμ|) for healthy individuals (ctl.) and patients (CAP, sepsis, H1N1). Due to the fluctuation-response relationship a magnitude of mean gene expression response depends on its variance. We estimated common variances for genes in healthy and CAP patients (top), healthy and sepsis patients (mid) and healthy and H1N1 patients (bottom). **C)** Correlations between variances in whole blood gene expression (σ^2^) and absolute changes in inter-individual gene expression variability (|Δσ^2^|) for control individuals (ctl.) and patients (CAP, sepsis, H1N1). **D)** Correlations between absolute changes in mean gene expression (|Δμ|) and in inter-individual gene expression variability (|Δσ^2^|).

Because of the fluctuation-response relationship [18], absolute changes in the mean log gene expressions (|Δμ|) in response to infection (CAP, H1N1) and sepsis correlated significantly with the variances of the log gene expressions (Figure 1B). Interestingly, we also noted significant correlations between the absolute changes in inter-individual gene expression variabilities (|Δσ^2^|) and the variances of log genes expressions (Figure 1C). Consequently, |Δμ| and |Δσ^2^| were also correlated (Figure 1D). Thus, we conclude that H1N1, CAP and sepsis result in coordinated changes in both the mean and heterogeneity of the expression of genes and that magnitudes of these changes depend on genes’ biological coefficients of variation.

### Ensemble gene noise response to infection and sepsis

Both the mean and variance relate to population (inter-individual) statistics reflecting distinct aspects of gene regulation. Changes in means fit the classical DGE view on gene response to a pathology and other biological processes, while changes in variances yield a view on heterogeneity of gene response. However, as we noted before (Figure 1 and S1), statistical inference of these changes is biased towards higher a significance for genes with a high biological coefficient of variation. Although changes in RNA copy number can serve in practical applications for diagnostics of a disease and clinical outcomes, inter-individual variability cannot be used for diagnosis. At the same time, stochastic fluctuations in gene expression remain attractive for the dissection of novel molecular mechanisms of a pathology. Therefore, we expect that estimation of ensemble gene noise may provide additional benefits for diagnostics by quantifying fluctuations, while being informative for personalized treatment.

We define ensemble gene noise as the variance of log-transformed, normalized expression levels for a collection of genes *G* = (*g*_1_,…, *g*_*i*_) encoding for either proteins of a pathway or subunits of a protein complex. To this end, we mapped genes to the KEGG-annotated pathways and the CORUM-annotated protein complexes [22, 23]. From the law of total variance: Var[*G*] = E[Var[*G*|*g*]]+ Var[E[*G*|*g*]], ensemble gene noise depends on the variability in expression of genes in ensemble (E[Var[*G*|*g*]]) and on their stoichiometry (Var[E[*G*|*g*]]) (Figure S1). Thus, the simple estimation of the variances (Var[*G*]) of gene ensembles for each individual might reflect alterations in function of biological pathways and protein complexes on the level of stoichiometry and gene noise.

We, then, correlated Var[*G*] for ensembles with H1N1, CAP and sepsis disease states. For H1N1 viral infection, disease state can be clearly ranked: non-infected (healthy) < early phase < late phase of infection, thus it represents an ordinal variable [28]. For CAP and sepsis patients, we assumed that a condition of the deceased patients was worse than that of the survived. We considered that healthy < survived < deceased can also be represented as ordinal disease state variable. Circumstantially, this is supported by distinct blood gene expression endotypes [8] and an increased gene expression heterogeneity (Figure 1A). Kendall rank correlation identified a number of pathways and protein complexes for which ensemble gene noise was positively and significantly associated with the disease state in H1N1 (FDR ≤ 0.05), and CAP and sepsis patients (Bonferroni-adjusted p ≤ 0.05) (Figure 2A). None of the pathways or gene complexes were negatively associated with the disease state at the specified significance thresholds. We used different p value adjustment procedures (FDR – less conservative, and Bonferroni – more conservative) for H1N1, CAP and sepsis patents due to the large differences in sample sizes (number of patients) between these data sets.

**Figure 2.**
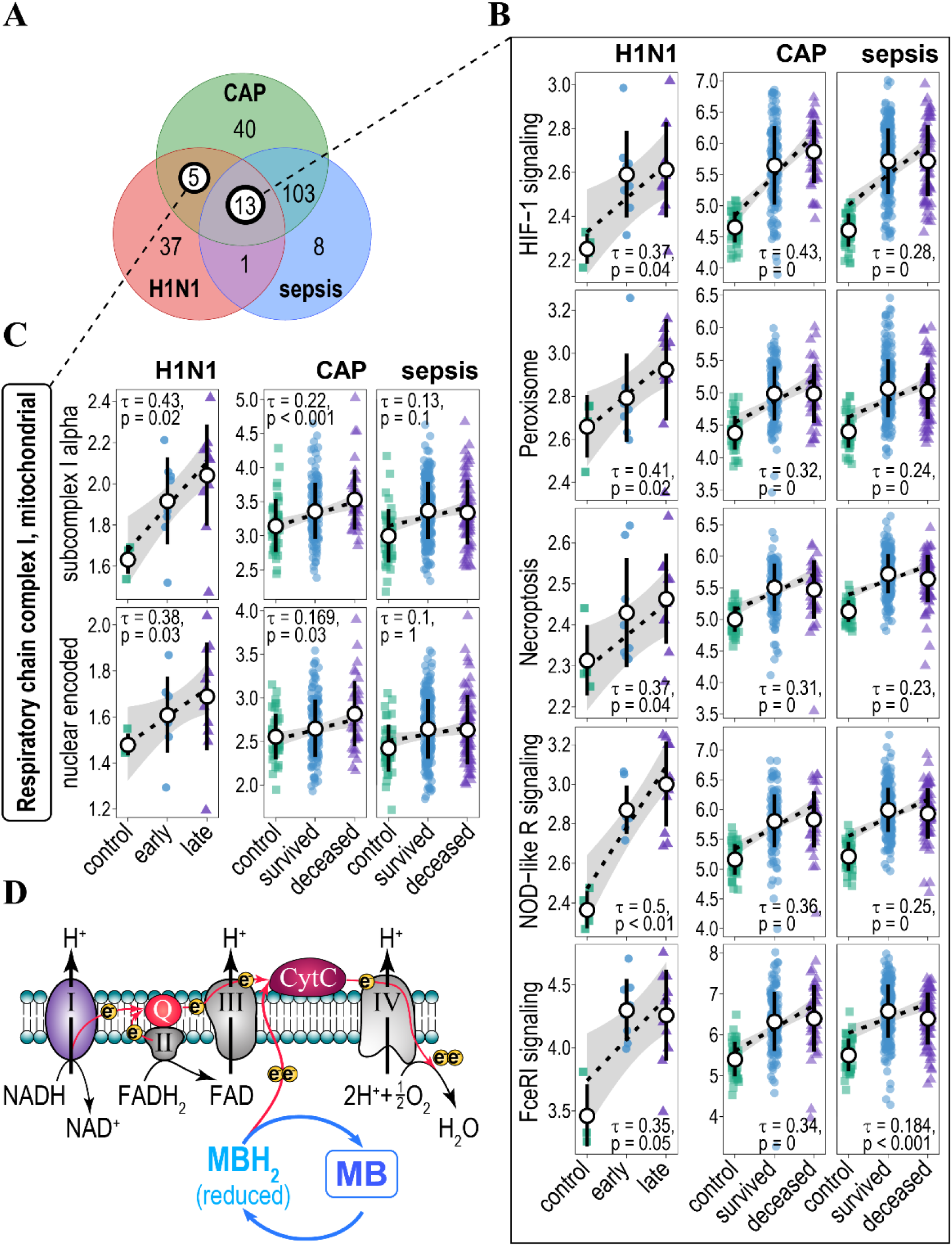
Association of ensemble gene noise with H1N1, CAP and sepsis disease states. **A)** Venn diagram of KEGG- and CORUM-annotated biological pathways/protein complexes for which ensemble gene noise associates positively (increases) and significantly with disease state. **B)** Plots of ensemble gene noise for genes involved in HIF-1 signalling, peroxisome, necroptosis, NOD-like receptor and Fc epsilon RI signalling pathways. Pathways were annotated by KEGG. Kendall tau, and FDR- (H1N1 patients) and Bonferroni-(CAP and sepsis patients) adjusted p-values are indicated. Rank-based regression trend lines and 95% confidence bands of ensemble gene noise association with the state of disease are shown. Black circles and whiskers indicate means and standard deviations. **C)** Plots of ensemble gene noise for genes encoding CORUM-annotated subunits of mitochondrial respiratory chain complex I (subcomplex I alpha – top panel and nuclear encoded subunits – bottom panel). Rank-based regression trend lines and 95% confidence bands of ensemble gene noise association with the state of disease are shown. **D)** Methylene Blue (MB) acts as an alternative electron donor to the electron transport chain (red arrows) by shuttling between redox states (MB – MBH_2_) and, thus, bypassing respiratory chain complex I. Respiratory chain complex I-IV and their substrates are indicated, Q – coenzyme Q10, CytC – cytochrome C. Electrons are indicated as yellow circles.

Out of all gene ensembles, 13 of them proved to be consistent and correlated to the increased disease state in ensemble gene noise in all three disease conditions (Figure 2A, B, Table S1A). Most of these gene ensembles (pathways) are known to be involved in the pathology of sepsis through multiple experimental evidences (Table 1), thus substantiating a power of ensemble gene noise analysis. However, ensemble gene noise yields novel insights into the molecular mechanisms of sepsis (H1N1, CAP or other-causes of sepsis) by suggesting a holistic mis-regulation in stoichiometry and gene noise for these gene ensembles.

**Table 1.** Role in sepsis of the pathways for which ensemble gene noise associates positively (increases) with the disease states (healthy < early/survived < late/deceased)

| Pathways/Complexes | Role in sepsis pathology | Reference |
| --- | --- | --- |
| KEGG: HIF-1 signalling pathway | Metabolic reprogramming of innate immune cells during the hyperinflammatory and immunotolerant phases of sepsis. | [51, 52] |
| KEGG: Peroxisome | Defective peroxisome recycling alters cellular redox homeostasis and leads to exaggerated oxidative stress response to endotoxin (infection) and sepsis. | [43] |
| KEGG: Necroptosis | Necroptosis is implicated in pulmonary diseases and sepsis-associated organ injury. | [53, 54] |
| KEGG: NOD-like receptor signalling pathway | Activation of Toll-like and NOD-like receptor signalling protects mice from polymicrobial sepsis-associated lethality. | [55] |
| KEGG: Fc epsilon RI signalling pathway | Fc receptors bind to antibodies attached to invading pathogens and their up-regulation can serve a potential biomarker for sepsis. Mice deficient for FCER1G gene encoding the γ-subunit of Fc epsilon RI show increased resistance to sepsis. | [56, 57] |
| KEGG: Autophagy - other | Autophagy is an adaptive protective process that eliminates damaged proteins, organelles and pathogens. It is thought to be a promising target in treatment of sepsis. | [58] |
| KEGG: Biosynthesis of amino acids | Sepsis results in significant disorders in amino acids metabolism. | [59] |
| KEGG: Glucagon signalling pathway | Glucagon levels negatively associate with clinical outcome in sepsis patients | [60] |
| KEGG: Propanoate (propionate) metabolism | Propionic acidemia caused by altered propionate metabolism often results in sepsis and death. | [61] |
| KEGG: Circadian rhythm | There is accumulating evidence for association between circadian misalignment and severity of inflammatory responses in sepsis. | [62] |
| KEGG: Dopaminergic synapse | Dopamine mediates neuroimmune communications and dopaminergic is implicated in inflammation and sepsis. | [63, 64] |
| KEGG: Amyotrophic lateral sclerosis (ALS) | ALS patients often develop pulmonary insufficiency and have increased risk of sepsis. | [65] |
| CORUM: Respiratory chain complex I, mitochondrial | Mitochondrial dysfunction resulting in reduced respiratory chain complex I activity and low ATP levels is a whole mark for sepsis. | [30] |
| KEGG: Osteoclast differentiation | Mean expression of osteoclast differentiation genes is up-regulated in human septic shock. | [66] |
| KEGG: Tight junction | Sepsis disrupts intestinal barrier which leads to a multiple organ dysfunction syndrome and alters the expression of tight junction proteins. | [67] |

We also identified 5 gene ensembles for which ensemble gene noise was positively and significantly correlated with the disease state in H1N1 and CAP patients (Figure 2A, Table S1B). However, ensemble gene noise for these pathways was also significantly increased in sepsis patients (t-test, Bonferroni adjusted p < 0.01) despite insignificant rank correlation. To that, some of these pathways can be implicated in the pathology of sepsis (Table 1). Two of these ensembles were represented by genes encoding mitochondrial respiratory chain complex I (Complex I) (Figure 2C). From the point of view of ensemble gene noise this suggests an altered stoichiometry and gene noise in the expression of the subunits of the Complex I which, as a result, might lead to its improper assembly and function in H1N1, CAP and sepsis patients.

Indeed, it has been established that the activity of the Complex I is decreased and correlates with the severity of sepsis [30]. Complex I is the first set of enzymes of the respiratory chain and it is the entry point for most electrons into the electron transport chain [31]. Interestingly, however, in case of the Complex I inhibition or deregulation, methylene blue (MB) can bypass it by acting as alternative redox mediator in the electron transport chain, thus, restoring mitochondrial respiration [32, 33] (Figure 2D). MB is also considered to be a promising therapeutic in treatment of septic shock [34, 35]. Thus, ensemble gene noise might provide a simple yet powerful explanatory shortcut, from the expression of thousands of genes to the function of gene ensembles and possible pharmaceutical targets.

### Predicting clinical outcome for CAP and sepsis patients from the ensemble gene noise

Treatment of sepsis is challenging and mortality rates among sepsis patients are high. Yet, prediction of clinical outcome is also challenging due to heterogeneity in the pathology [8] and gene expression (Figure 1A). Recently, Molecular Diagnosis and Risk Stratification of Sepsis (MARS) consortium identified the Mars1 gene expression endotype which was significantly associated with acute (28-day) mortality, however, for other endotypes Mars2-4 poorly discriminated between the survival and mortality of patients [8]. Thus, we wondered whether the clinical outcome (mortality) could be predicted from the ensemble gene noise.

To this end, we trained binary logistic gradient boosted regression tree models using survival and acute mortality as a binary response variable for clinical outcome and patients’ age and blood ensemble gene noise as models’ features. The models were trained with XGBoost [36]. The CAP and sepsis patients were split into discovery (263 patients: 105 CAP and 158 sepsis patients) and validation (216 patients: 78 CAP and 138 sepsis patients) cohorts following GSE65682 annotation [8]. Within the cohorts the mortality rates were 26.2% for CAP and sepsis patients (23.8% for CAP and 27.8% for sepsis patients) in the discovery cohort, and 20.8% for CAP and sepsis patients (19.2% for CAP and 21.7% for sepsis patients) in the validation cohort.

Overall, class-imbalance, noise due to the inter-individual heterogeneity and high-dimensionality of model features are among the major problems of machine learning [37]. In part, ensemble gene noise leads to a reduction in inter-individual variability (Figure S3) and in dimensionality as model features are represented not by individual genes, but by collections of genes. Nonetheless, we further reduced the number of ensemble gene noise features in models by t-test feature selection. For this, we compared gene ensembles noise between survived and deceased patients in the discovery cohorts. The p-value cut-offs for the model features were selected based on maximization of models’ training accuracy (see Methods). XGBoost hyper-tuning parameters: learning rate, complexity, depth, *etc.* were optimized based on the cross-validation. To avoid overfitting, we used early epoch stopping, which was estimated from the test fold of the discovery cohort (see Methods). Because of the class-imbalance, AUC (area under the receiver operating characteristic (ROC) curves) was used to evaluate the model performance. The validation cohorts were hidden from the feature selection and training.

Figures 3A and 3B show model scores and ROC curves for the model, predicting mortality/survival for the CAP and sepsis patients in the discovery and validation cohorts. AUCs for the discovery and validation cohorts were 0.871 and 0.707 respectively, suggesting a reasonable accuracy of the model. However, from the model scores, and evaluation of the model specificity/sensitivity it appears that the model is biased towards the prediction of major class (survived) (Figure 3A, Table 2 and Table S2A). Thus, class prediction balanced accuracies (bACC = Specificity/2 + Sensitivity/2) were 0.799 and 0.701 for the discovery and validation cohorts respectively. Nonetheless, the survival probability for patients predicted to have a high risk of mortality was significantly lower than the survival probability of patients predicted to have low risk of mortality in both discovery and validation cohorts. To that, our model better predicts the risks of mortality as compared to the Mars1 endotype inferred from the log gene expression unsupervised learning (Figure 3C) [8]. Potentially, this could be due to a lower inter-individual variability of gene ensembles noise as compared to log gene expression (Figure S3).

**Table 2.** Prediction accuracy of the models.

| **Metric** | **CAP/sepsis patients** |  | **CAP patients** |  | **Sepsis patients** |  |
| --- | --- | --- | --- | --- | --- | --- |
|  | **discovery** | **validation** | **discovery** | **validation** | **discovery** | **validation** |
| **bACC** | 0.799 | 0.701 | 0.802 | 0.798 | 0.779 | 0.761 |
| **Sensitivity** | 0.754 | 0.6 | 0.88 | 0.867 | 0.75 | 0.8 |
| **Specificity** | 0.845 | 0.801 | 0.725 | 0.73 | 0.807 | 0.722 |

**Figure 3.**
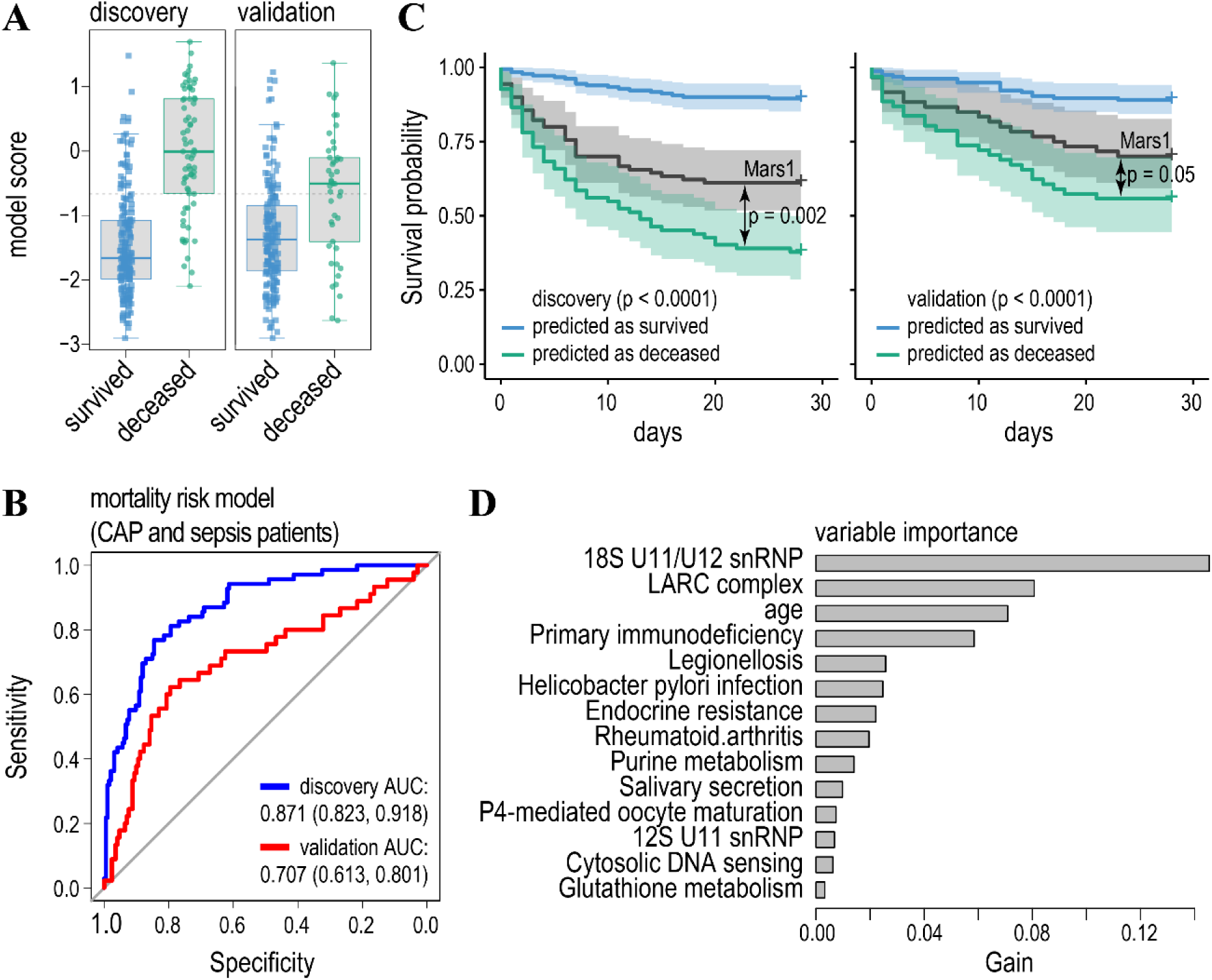
Model predicting mortality/survival of CAP and sepsis patients. **A)** Boxplots of the model scores predicting mortality/survivorship in the discovery (left) and validation (right) cohorts. The model was trained on the same as published discovery cohort by the gradient boosted regression tree and validated on independent cohort[8]. Dashed lines indicate threshold levels of classification. The threshold was calculated by maximizing a product of the specificity and sensitivity of the model prediction in the discovery cohort. Further details of model accuracy are given in Tables 2 and S2. **B)** Receiver operating characteristic curves (ROC) for the model predicting mortality (end point – survival or death within 28 days after treatment) in CAP and sepsis patients (blue line – discovery cohort, red line – validation cohort). Features were selected by the t-test comparing ensemble gene noise between the survived and deceased patients in the discovery cohort to achieve maximum prediction accuracy for the discovery cohort. Values for the area under the ROC curve (AUC) are indicated. **C)** Survival probability for the patients predicted to have low (blue line) and high (green line) risk of mortality for the discovery (left panel) and validation (right panel) cohorts. p-values indicate significant differences in hazards for the predicted classes (survival/mortality) according to the Cox proportional-hazards model. Black lines - survival probability of patients with Mars1 endotype [8] was compared with the predicted deceased class for the discovery and validation cohorts. **D)** Variable importance of the model ranks ensemble gene noise features according to their relative contribution (gain).

In an attempt to increase the prediction accuracy, we trained to separated gradient boosted tree models for CAP (Figure 4) and sepsis (Figure 5) patients. Indeed, in both cases the accuracy of the prediction of the minor class (deceased patients) increased (Table 2 and S2) in both discovery and validation cohorts. Likewise, AUCs for the validation cohorts were also higher as compared to the model predicting mortality for both (CAP and sepsis) type of patients (compare Figure 4B and 5B with Figure 3B). To that, differences in AUCs between discovery and validation cohorts were lower for the models predicting mortality separately for CAP and sepsis patients as for the model trained on both type of patients. This was especially evident for the model predicting mortality for the CAP patients (Figure 4B). Thus, knowing the cause of sepsis improves the prediction accuracy of the models.

**Figure 4.**
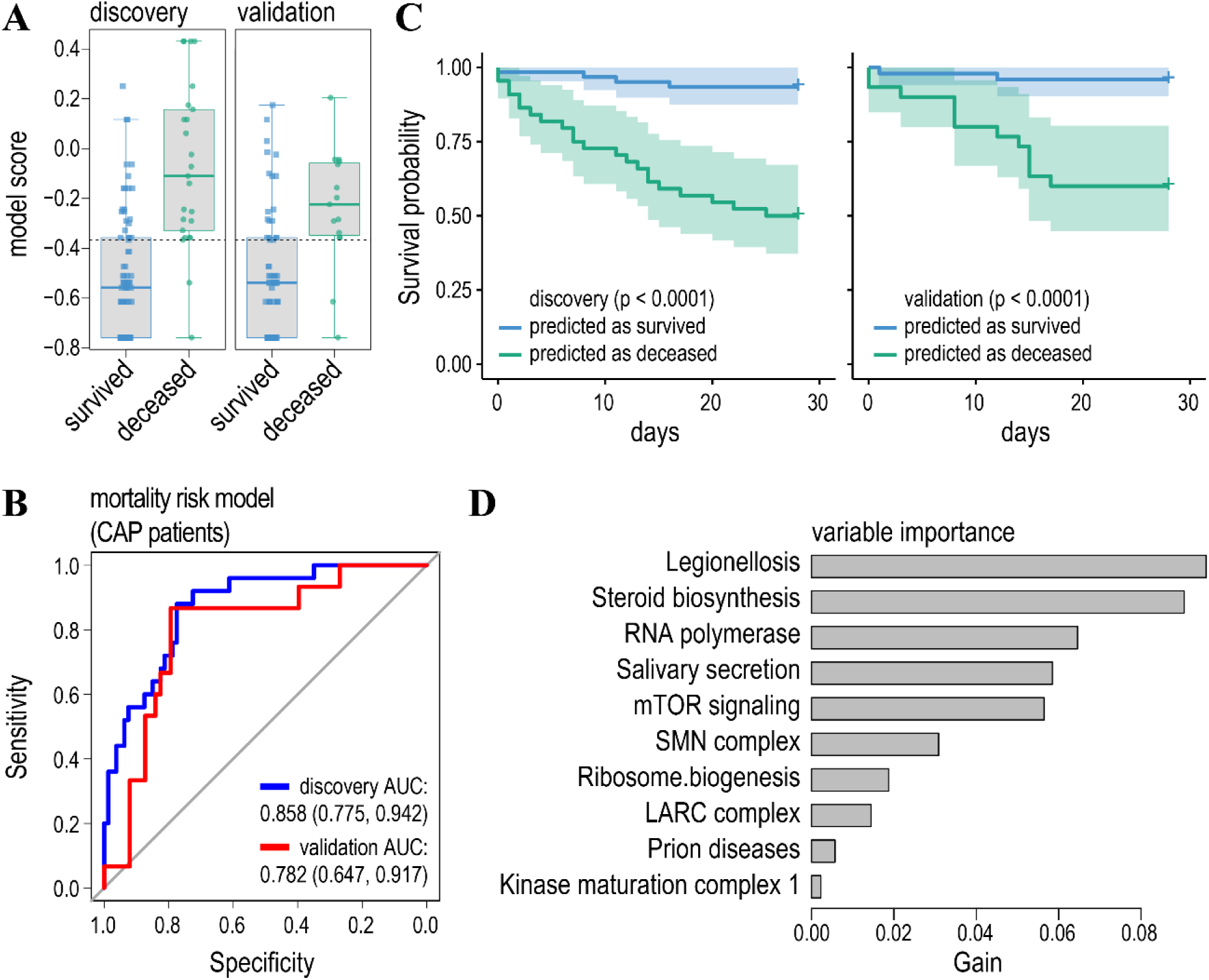
Model predicting mortality/survival of CAP patients. **A)** Boxplots of the model scores predicting mortality/survivorship in the discovery (left) and validation (right) cohorts. **B)** ROC curves for the model predicting mortality in CAP patients in the discovery (blue line) and validation (red line) cohorts. Cohorts were partitioned as in [8]. **C)** Survival probability for the patients predicted to have low (blue line) and high (green line) risk of mortality for the discovery (left panel) and validation (right panel) cohorts. **D)** Relative contribution of ensemble gene noise features to the model.

**Figure 5.**
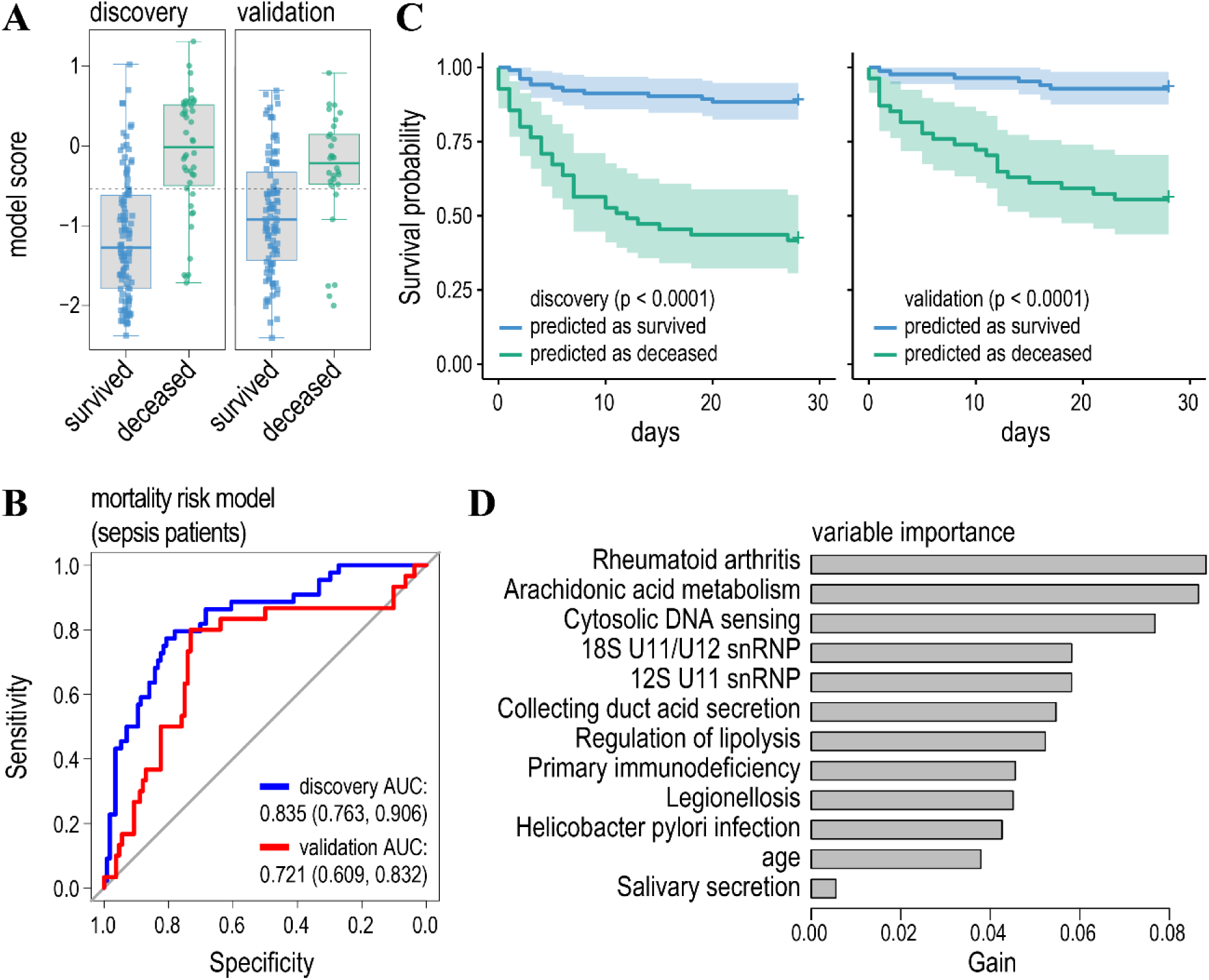
Model predicting mortality/survival of sepsis patients. **A)** Boxplots of the model scores. **B)** ROC curves for the model predicting mortality in sepsis patients. **C)** Survival probability for the patients predicted to have low (blue line) and high (green line) risk of mortality. **D)** Relative contribution of ensemble gene noise features to the model.

Finally, it has to be noted that both the feature selection and gradient boosted regression trees allow for the ranking of the model features’ importance (Figures 3D, 4D, 5D and Table S3). First, it turned out that a patients’ age does not noticeably contribute to the prediction of mortality in CAP patients and it ranks low in the prediction of mortality of sepsis patients. Second, high ranking gene ensembles (pathways) could be immediately associated with host response to infection and, thus, pathology of the sepsis. These include legionellosis (a pathway responsible for atypical pneumonia caused by *Legionella* bacteria), epithelial cell signalling in *Helicobacter pylori* infection and leishmaniasis, and imbalances in these pathways either caused by corresponding infections or immune activation could lead to the sepsis [38-40]. To that, ensemble gene noise in immune pathways, such as rheumatoid arthritis and primary immunodeficiency, contribute to the prediction of clinical outcome in sepsis patients (Figure 3D, 5D and Table S3). Thus, we conclude that the ensemble gene noise uncovers novel approaches and insights to the discovery of biomarkers, prediction of clinical outcome and to the molecular mechanisms of a pathology from the point of view of imbalances in stoichiometry and gene noise of expression in gene ensembles.

## Discussion

Here we attempted a dissection of molecular mechanisms of human pathology, exemplified by H1N1 infection, CAP and sepsis, through a prism of ensemble gene noise. Unlike classical DGE, ensemble gene noise allows for the identification of imbalances in the expression of entire gene circuits, rather than individual genes on the level of stoichiometry and gene noise. This approach offers an alternative, but non-mutually exclusive to the DGE interpretation of a molecular basis of disease and both have their own strengths and weaknesses.

We noted in the introduction that due to a fluctuation-response a statistical inference of DGE might be biased towards genes with a high inter-individual variability, *i.e.* “noisy” genes (Figure S1) [18, 19]. However, the same applies to ensemble gene noise (Figure S4). This imposes a certain problem to the interpretation of both DGE and ensemble gene noise. On one hand, it can be suggested that large deviations in expression of genes and ensembles, which are naturally prone to high fluctuations, might not be causative for a disease, as an organism is already adapted to such variations. On the other hand, these genes/ensembles themselves might play an important adaptive role [41] and their over-response could lead to a disease. At the moment it seems difficult to come to a resolution between these two possibilities, but they should be considered, specifically in identification of pharmaceutical targets: genes or gene ensembles (pathways, protein complexes).

As compared to DGE, ensemble gene noise provides a holistic interpretation to mis-regulation in gene expression under pathologic or other conditions. As it operates on the level of gene ensembles it does not require gene set enrichment analysis (GSEA), thus it circumvents potential pitfalls of GSEA associated with the cut-off problem of DGE [24, 25]. As any gene expression analysis ensemble gene noise relies on the quality and completeness of pathways and the protein complexes’ annotation. Finally, we noted that inter-individual variability of ensemble gene noise is significantly less than that of individual gene expression (Figure S3). This, in turn, might improve the accuracy of diagnostic and clinical outcome models. Though it might come at the expense of less features being available for the selection and training of models. At the same time, in future studies, both DGE and ensemble gene noise could be combined.

In this study we applied the concept of ensemble gene noise to the analysis of critically ill H1N1, CAP and sepsis patients [8, 28]. We noted a large-scale gene response in two dimensions: on the level of mean gene expression and on the level of variance (inter-individual variability). Interestingly, both responses were correlated (Figure 1D) and both were dependent on gene variance suggesting that the fluctuation-response might drive changes in these two parameters of gene expression co-ordinately [18]. In all three cases (H1N1, CAP and sepsis), inter-individual variability was increased for a bulk of the genes. Consequently, we only identified pathways or gene complexes for which ensemble gene noise was significantly increased for H1N1, CAP and sepsis patients as compared to healthy individuals. This suggests that inter-individual gene expression variability is a prominent driver of ensemble gene noise in these patients.

Because viral/bacterial infections and sepsis result in overwhelming gene expression response, it is difficult to identify a reasonably small set of either genes or gene ensembles for biological interpretation. Thus, we only focused on the pathways (protein complexes) for which ensemble gene noise increased in all three cases and correlated these with a disease state (Figure 2A). From this intersection we inferred 13 pathways most of which have been previously implicated in sepsis (Table 1). To that, 5 pathways (protein complexes) showed significant association of ensemble gene noise with H1N1 infection phase and CAP disease state and for which ensemble gene noise also increased significantly in sepsis patients (Figure 2A). Potentially, these pathways could be targeted for adjuvant treatment of sepsis. Especially, we consider mitochondrial respiratory chain complex I (Complex I) (Figure 2D) and peroxisome promising for pharmaceutical targeting. Increased ensemble gene noise for the Complex I would imply either altered stoichiometry, or increased gene expression noise for genes encoding subunits of the Complex I or both. As a result, this might lead to improper assembly of the Complex I and affecting its function. The impaired Complex I function can be bypassed by an alternative redox mediator, such as methylene blue [32, 33]. To that, methylene blue is a selective inhibitor of the nitric oxide–cyclic guanosine monophosphate (NO–cGMP) pathway [35] and increased NO levels is a hallmark of sepsis [42]. Some clinical studies have already indicated a beneficial role of methylene blue in the treatment of sepsis [34, 35]. Similar to mitochondrial respiration, peroxisomes also play an important role in the pathology of sepsis as the dysfunction of peroxisomes results in oxidative stress [43]. Again, an increased ensemble gene noise for peroxisome pathway indicates a potential mechanism for such dysfunction in H1N1, CAP and sepsis patients. Potentially peroxisome biogenesis could be restored by 4-phenylbutyrate and there several studies indicating its positive role in treatment of sepsis [44, 45]. Considering future directions, it could be proposed that search for epigenetic modulators of ensemble gene noise might represent a novel pharmaceutical avenue for adjuvant treatments of sepsis.

Finally, we explored the possibility to use ensemble gene noise in the prediction of clinical outcomes. Previously some promising biomarkers and gene expression endotypes associated with septic shock and mortality have been identified based on DGE analysis [8, 9]. However, as already mentioned, ensemble gene noise looks at gene expression from a different, yet complementary, angle, thus enabling the identification of novel pathways and biomarkers for sepsis and other diseases. To that, models predicting pathology based on ensemble gene noise could potentially be more robust, as inter-individual variability for ensemble gene noise is lower than that for log gene expression (Figure S3). Furthermore, Gradient boosted regression tree models trained on CAP and sepsis patients to predict their mortality had a good accuracy on validation cohort (Figure 3, Table 2). These outperformed predictions based on the Mars1 gene expression endotype, which was shown to associate with a poor prognosis [8], both on the discovery and validation cohorts (Figure 3C). Interestingly, some ensemble gene noise features selected statistically for the models predicting mortality in both CAP/sepsis-, CAP- and sepsis-patients couldimmediately be related to the host’s response to infection. For example, increases in ensemble gene noise in legionellosis, epithelial cell signalling in *Helicobacter pylori* infection and leishmaniasis pathways could potentially serve as biomarkers of sepsis and its outcome.

In conclusion, here we showed a potential of ensemble gene noise in the biological interpretation of a disease, the identification of pharmaceutically targetable pathways, novel biomarkers, and the prediction of clinical outcome. Together, we believe that ensemble gene noise analysis could be broadly applied alongside with DGE to dissect molecular mechanisms of the pathology in two complementary dimensions: in Jacob-Monod dimension of specific gene regulation and in a novel dimension of holistic gene circuit regulation.

## Methods

### Data resources and processing

GSE65682 Affymetrix Human Genome U219 Array whole blood gene expression profiles were used for the analysis of community/hospital acquired pneumonia (CAP) and sepsis patients [8, 11]. In brief, the cohort consisted of 42 healthy individuals (24 males, 18 females), 183 CAP patients (111 males, 72 females) and 296 sepsis patients (161 males, 135 females). The mean age of CAP (61.5±1.2) and sepsis (60.6±0.8) patients did not differ significantly (*t*(350.37) = 0.59, p = 0.56), however healthy individuals were significantly younger as compared to CAP (*t*(54.0) = 4.7, p < 0.001) and sepsis (*t*(47.2) = 4.6, p < 0.001) patients. Out of 183 CAP patients, 40 died within 28 days and out of 296 sepsis patients, 74 died within 28 days. Thus, we divided CAP and sepsis patients into survived and deceased groups, considering these two states as an ordered factor (ordinal) variable (survived < deceased).

GSE21802 Illumina human-6 v2.0 expression bead-chip whole blood gene expression profiles were used for the analysis of H1N1 infected patients [28]. The cohort consisted of 4 healthy individuals and 19 H1N1 patients (8 in early and 11 in late phase of the disease). The early phase was defined as early, from the onset of symptoms - day 0 to day 8, and late – from day 9 and above. The statistics of the cohorts is given in Table 1 of [28], however neither sex nor age assignments were available for the patients from the GSE21802 series annotation.

GSE65682 microarrays signal intensities were pre-processed (background corrected and RMA-normalized) with the Bioconductor *oligo* package [46]. Lowly-expressed and outlier genes were identified in high dimensions using the spatial signs (*sign2*) algorithm of *mvouliter* R package with a critical value for outlier detection at 0.9. The robust principal components explained a variance of 0.95 [47]. GSE21802 signal intensities significantly above the background were quantile normalized [48]. Genes were annotated with Bioconductor *hgu219.db* and *illuminaHumanv2.db* database packages for GSE65682 (8826 genes) and GSE21802 (7240 genes) respectively.

### Statistical analysis of gene expression variability and ensemble gene noise

Statistical analysis was done using R and R/Bioconductor packages [49].

To estimate the inter-individual gene expression variability for healthy, CAP and sepsis patients we accounted for age as a random effect. To this end, we used Generalized Additive Model for Location, Scale and Shape (GAMLSS) [20, 29]. In brief, for normally distributed log-transformed microarray intensities (*Y* = *log*(*X*), *Y* ∼*N*(*μ*_y_, *σ*_y_)), GAMLSS allows for the modelling of both parameters of gene expression (mean and dispersion):

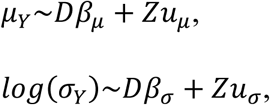

where 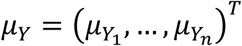 and 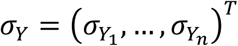 are the vectors of means and dispersions for *Y* = (*Y*_1_,…, *Y*_n_). *D* - *n* x *p* fixed effect design matrix for the disease state (healthy, survived, deceased). 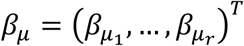 and 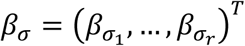 – estimated fixed effect coefficients for mean and dispersion. *Z*-*n* x *k* random effect design matrix for age (age was binned into 10 deciles). 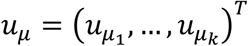 and 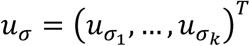 – estimated random effect coefficients for mean and dispersion, where *u*∼*N*(0, *δ*). With GAMLSS it is also straightforward to test for the significance of a factor effect on either the mean, the variance, or both with likelihood ratio test [20].

Gene ensemble lists were generated by the mapping of genes to the KEGG-annotated biological pathways or CORUM-annotated subunits of mammalian protein complexes [22, 23]. Their gene noise was estimated for each individual by calculating the variances of log-transformed expressions of genes for each ensemble (Figure 1S). Estimates of gene ensembles noise were correlated with the disease states (healthy < early phase < late phase for H1N1 and healthy < survived < deceased for CAP and sepsis) by Kendall rank correlation, treating the disease state as an ordinal variable. Linear trends between disease states and ensembles gene noise were estimated by rank-based regression [50].

### Gradient boosted regression tree models

To predict the mortality of CAP and sepsis patients we trained gradient boosted regression tree models with a scalable tree boosting system XGBoost [36] using mortality within 28 days as a binary response variable, and ensemble gene noise and age as independent model features. To this end, we split individuals into discovery and validation cohorts following exactly the same partitioning as annotated in GSE65682 [8]. Then, we trained 3 models: 1) a model predicting mortality for CAP and sepsis patients, 2) a model predicting mortality for CAP patients, and 3) a model predicting mortality for sepsis patients. Models features were preselected using discovery cohorts by *t* test comparing ensembles gene noise for survived and deceased patients to maximize the accuracy of XGBoost training on the discovery data sets. For CAP and sepsis (1), and sepsis (3) models, the cut-off for the model features was set at p ≤ 0.01, and for the CAP model (2) – at p ≤ 0.05. The XGBoost hyper tuning parameters (learning rate (η), complexity (γ), depth, *etc.*) were optimized by cross validation. To avoid overfitting, we found early epoch stopping parameters by randomly splitting of the discovery cohort into two equal folds: training and test. Then, the validation cohorts, which were hidden from feature selection and model training, were used to verify the accuracy of the final models.

## Acknowledgements

We thank Oleg Derkatch who motivated this work and Dr. Laurent Schwartz, Olga and Errol Fontanellaz who pointed to a potential use of methylene blue. This work has been supported by the Russian Science Foundation grant: 20-14-00055 to YMM and Gene Learning Association.

## Supplementary material

**Figure S1.**
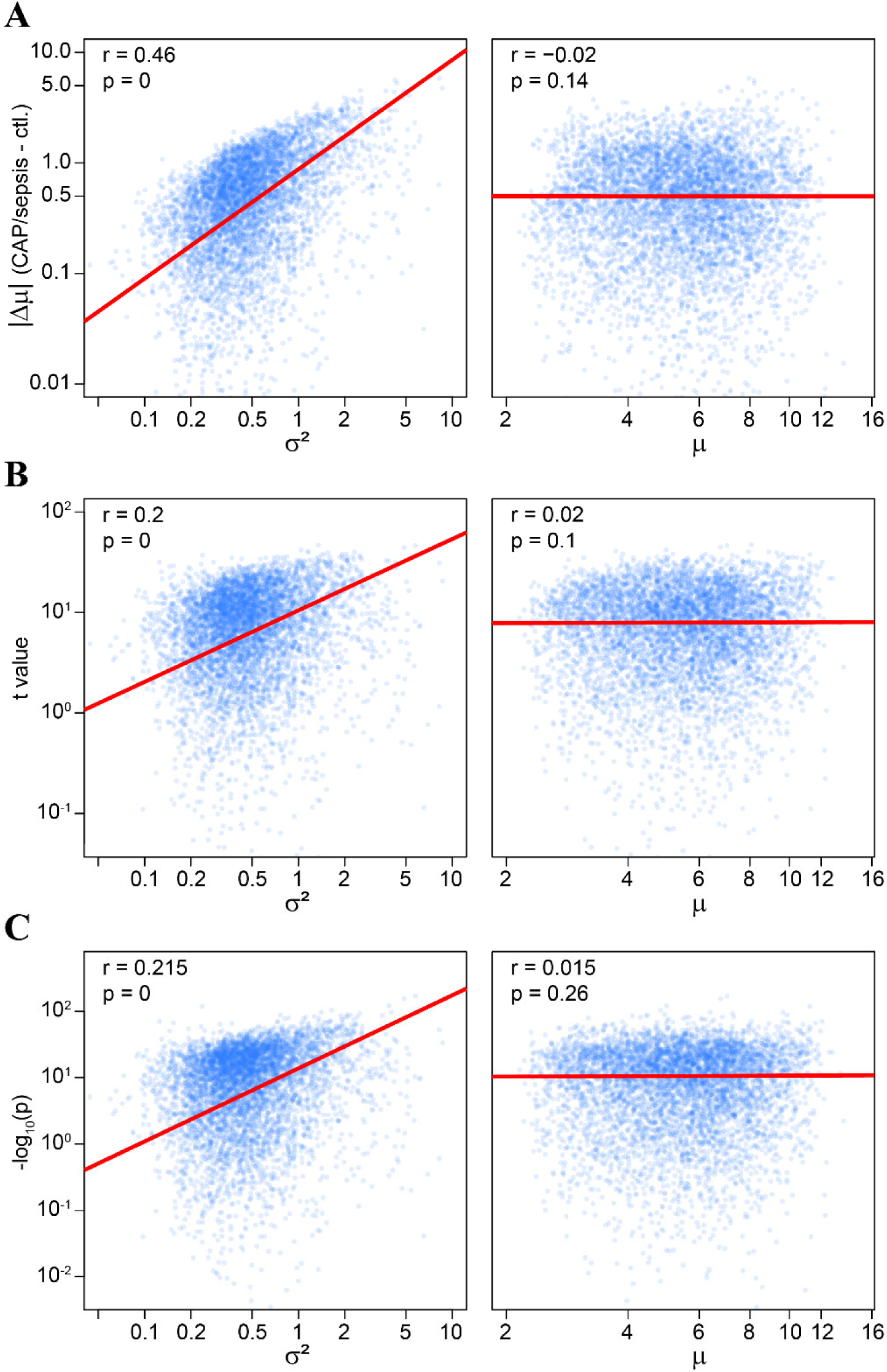
Fluctuation-response relation biases statistical inference of DGE. **A)** The difference of means of log-transformed bell-shaped gene expression values (*Y* = *log*(*X*)) are proportional to the variance or the squared coefficient of variation (*cv*^*2*^) of untransformed variable (*X*): 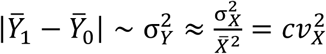. Considering RNA copy number (*X*) to be mixed-Poisson (negative-binomial as a specific case) random variable, for large 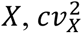 approximates overdispersion parameter or biological coefficient of variation (*bcv*^*2*^) [20]. A scatterplot on the left panel illustrates fluctuation-response relation (a correlation between absolute differences and variances of log-transformed expression values) for whole-blood gene expression profiles of healthy (ctl.) individuals and CAP pneumonia/sepsis patients. The data has been taken from [8]. This relation is monotonic, but non-linear, suggesting a deviation from linear coupling between fluctuation (pneumonia/sepsis) and gene expression response. There is no correlation between differences and means of log-transformed expression values (right panel). **B-C)** In the presence of fluctuation-response relation statistical inference will be biased. For example, Student’s *t*-test often used to assess DGE will relate positively to a variance of log-transformed gene expression as: 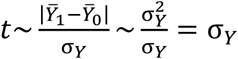. A scatterplot on the left panel shows correlation between Student’s *t* statistic and variance (**B**) and Bonferroni-adjusted p values and variance (**C**). There is no correlation of *t* and *p* values with means of log-transformed expression values (right panel).

**Figure S2.**
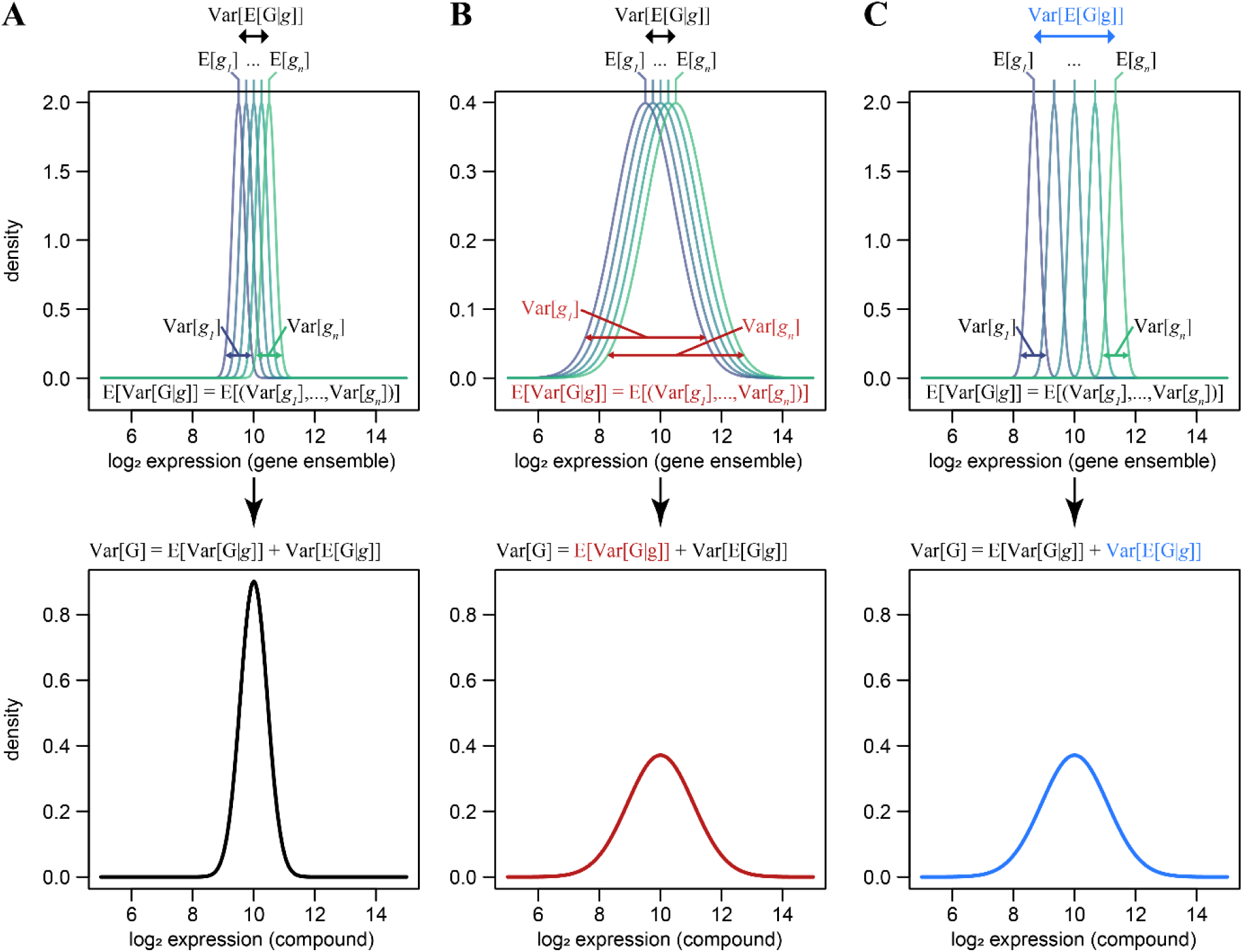
Ensemble gene noise. **A)** Let *G* be a collection of genes (*g*_1_,…, *g*_*n*_) belonging to either a biological pathway or encoding subunits of a protein complex. Then, from the law of total variance Var[*G*] = E[Var[*G*|*g*]] + Var[E[*G*|*g*]], *i.e.* ensemble gene noise (Var[*G*]) sums from the expected value of genes’ variances (E[Var[*G*|*g*]]) and the variance in genes’ mean expression (Var[E[*G*|*g*]]). The top panel illustrates hypothetical distributions of expressions of genes in ensemble (*g*_*i*_), the bottom panel is derived distribution of gene ensemble (*G* = (*g*_1_,…, *g*_*n*_)). **B, C)** Top panel, changes in variances (**B**) and/or expectations (**C**) of genes expression will eventually change ensemble gene noise (bottom panel).

**Figure S3.**
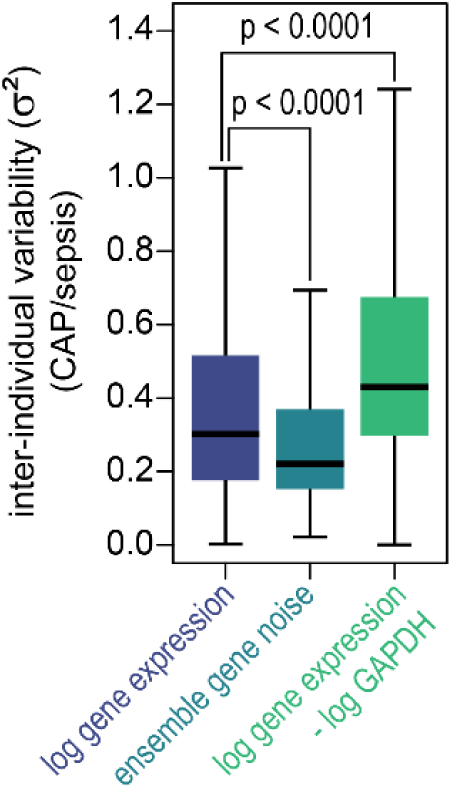
Comparison of inter-individual variability for log gene expression and ensemble gene noise for CAP and sepsis patients. Boxplots illustrating population variances for log gene expression, ensemble gene noise and log gene expression normalized to the GAPDH. Inter-individual variability is significantly less for ensemble gene noise as compared to the log gene expression (according to t-test) and it is higher for GAPDH normalized log gene expressions. The latter follows from the fact that Var[*log*(*X*) − *log*(GAPDH)] ≈ Var[*log*(*X*)] + Var[*log*(GAPDH)]. Thus, estimating DGE by PCR, which usually requires normalization to some housekeeping gene, results in increased inter-individual variability. Ensemble gene noise can be estimated from PCR without normalization to a reference gene.

**Figure S4.**
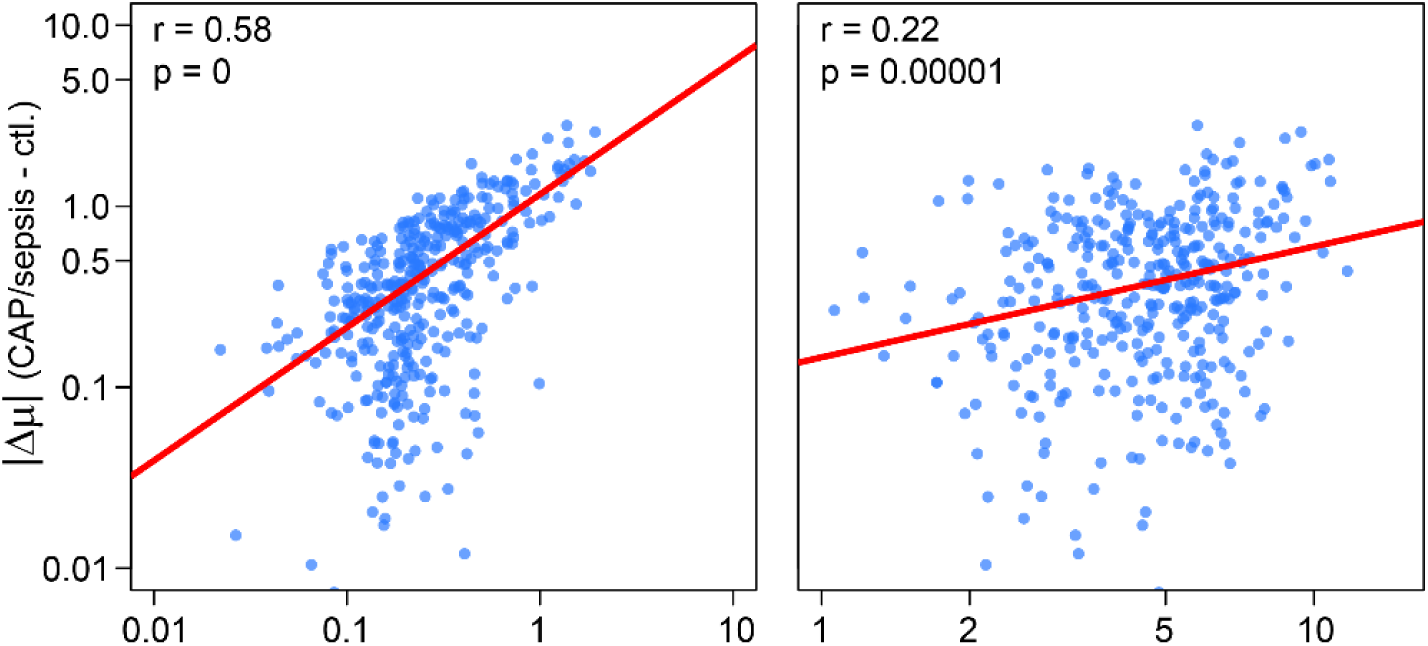
Fluctuation-response relation for ensemble gene noise. Differences of means of ensembles gene noise are proportional to the inter-individual variability in ensembles gene noise. A scatterplot on the left panel illustrates fluctuation-response relation (a correlation between absolute differences and variances of ensembles gene noise) for whole-blood gene expression profiles of healthy (ctl.) individuals and CAP pneumonia/sepsis patients. There is also a modest correlation between absolute changes in ensembles gene noise and the means (right panel).

**Table S1.**
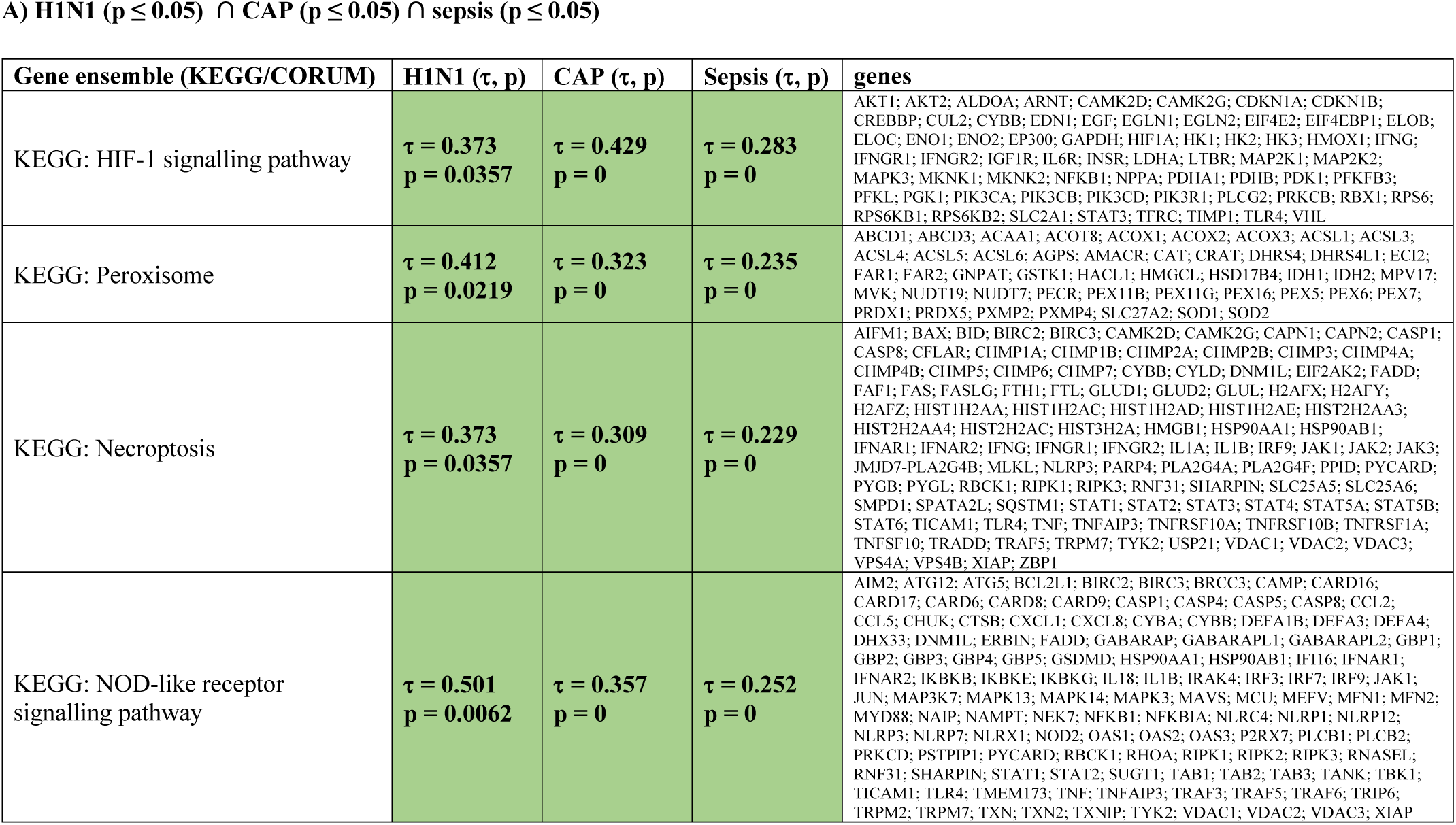
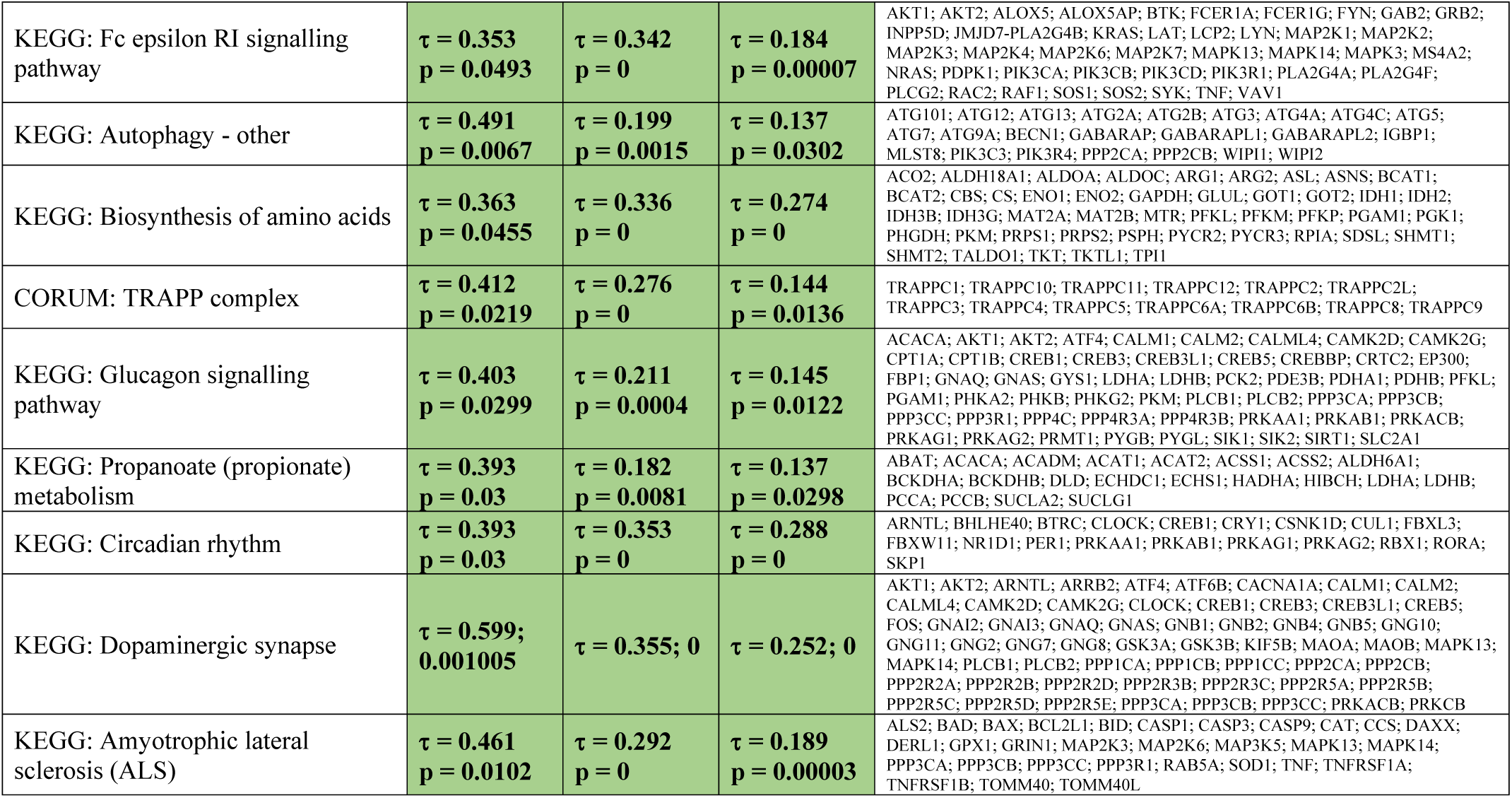
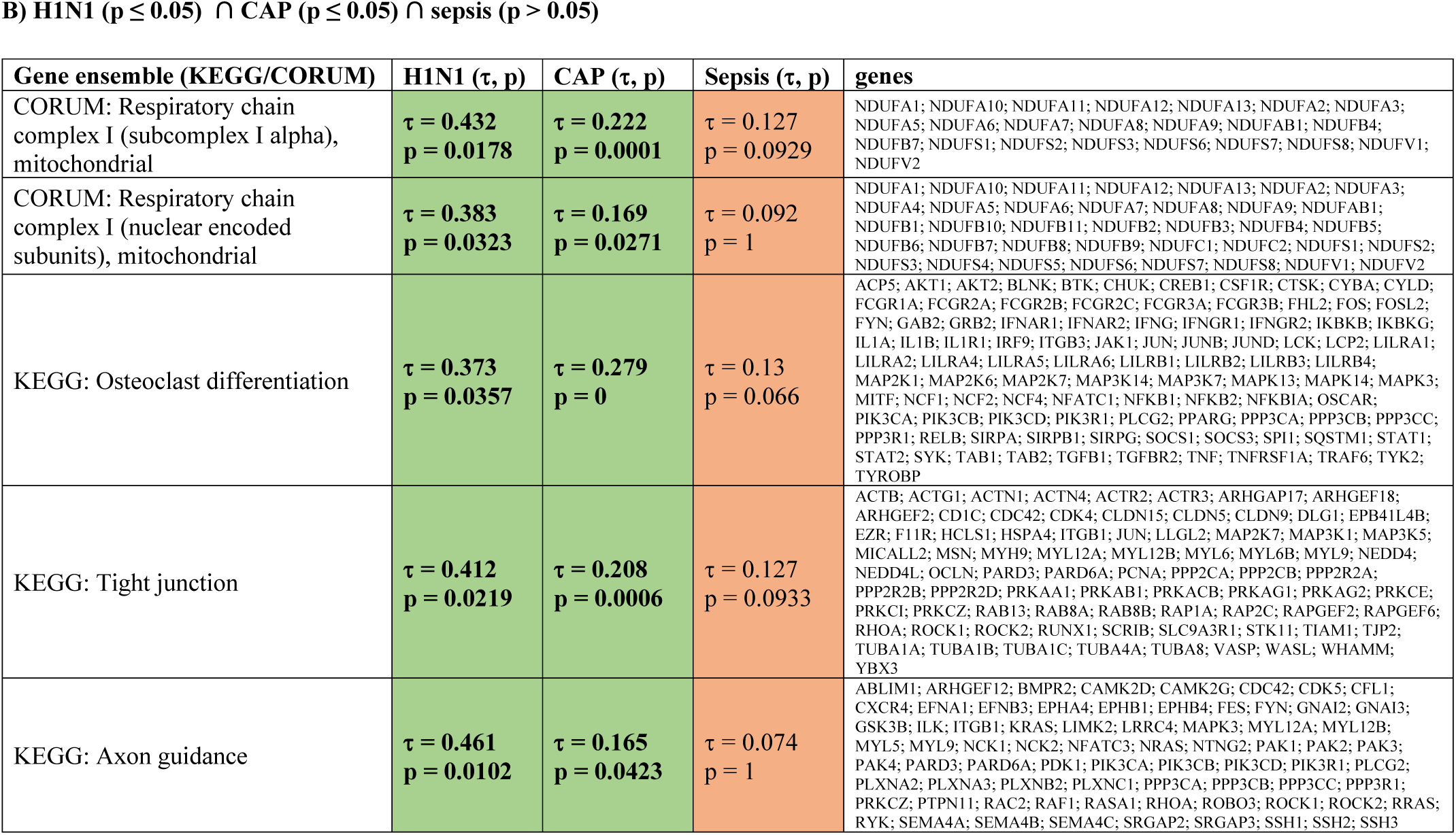
Kendall correlation of ensemble gene noise with H1N1, CAP and sepsis disease states.

**Table S2.** Confusion tables for the models predicting mortality/survivorship of CAP/sepsis patients

**A) CAP and sepsis patients**
| discovery |  | Actual |  |
| --- | --- | --- | --- |
|  |  | survived | deceased |
| predicted | survived | 164<br>(84.5%) | 17<br>(24.6%) |
|  | deceased | 30<br>(15.5%) | 52<br>(75.4%) |

| validation |  | Actual |  |
| --- | --- | --- | --- |
|  |  | survived | deceased |
| predicted | survived | 137<br>(80.1%) | 18<br>(40.0%) |
|  | deceased | 34<br>(19.9%) | 27<br>(60.0%) |

**B) CAP patients**
| discovery |  | Actual |  |
| --- | --- | --- | --- |
|  |  | survived | deceased |
| predicted | survived | 58<br>(72.5%) | 3<br>(12.0%) |
|  | deceased | 22<br>(27.5%) | 22<br>(88.0%) |

| validation |  | Actual |  |
| --- | --- | --- | --- |
|  |  | survived | deceased |
| predicted | survived | 46<br>(73.0%) | 2<br>(13.3%) |
|  | deceased | 17<br>(27.0%) | 13<br>(86.7%) |

**C) Sepsis patients**
| discovery |  | Actual |  |
| --- | --- | --- | --- |
|  |  | survived | deceased |
| predicted | survived | 92<br>(80.7%) | 11<br>(25.0%) |
|  | deceased | 22<br>(19.3%) | 33<br>(75.0%) |

| validation |  | Actual |  |
| --- | --- | --- | --- |
|  |  | survived | deceased |
| predicted | survived | 78<br>(72.2%) | 6<br>(20.0%) |
|  | deceased | 30<br>(27.8%) | 24<br>(80.0%) |

**Table S3.**
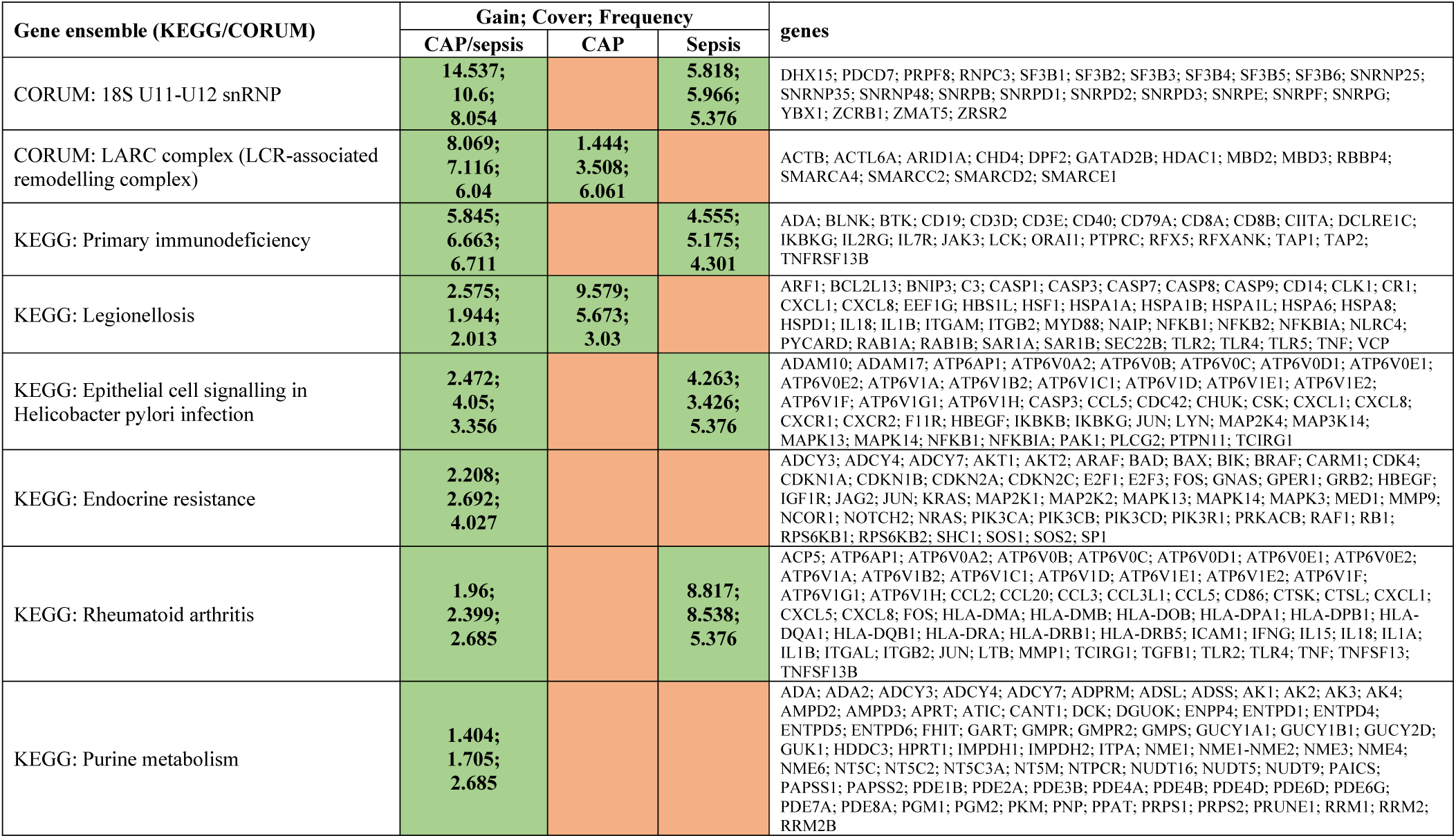
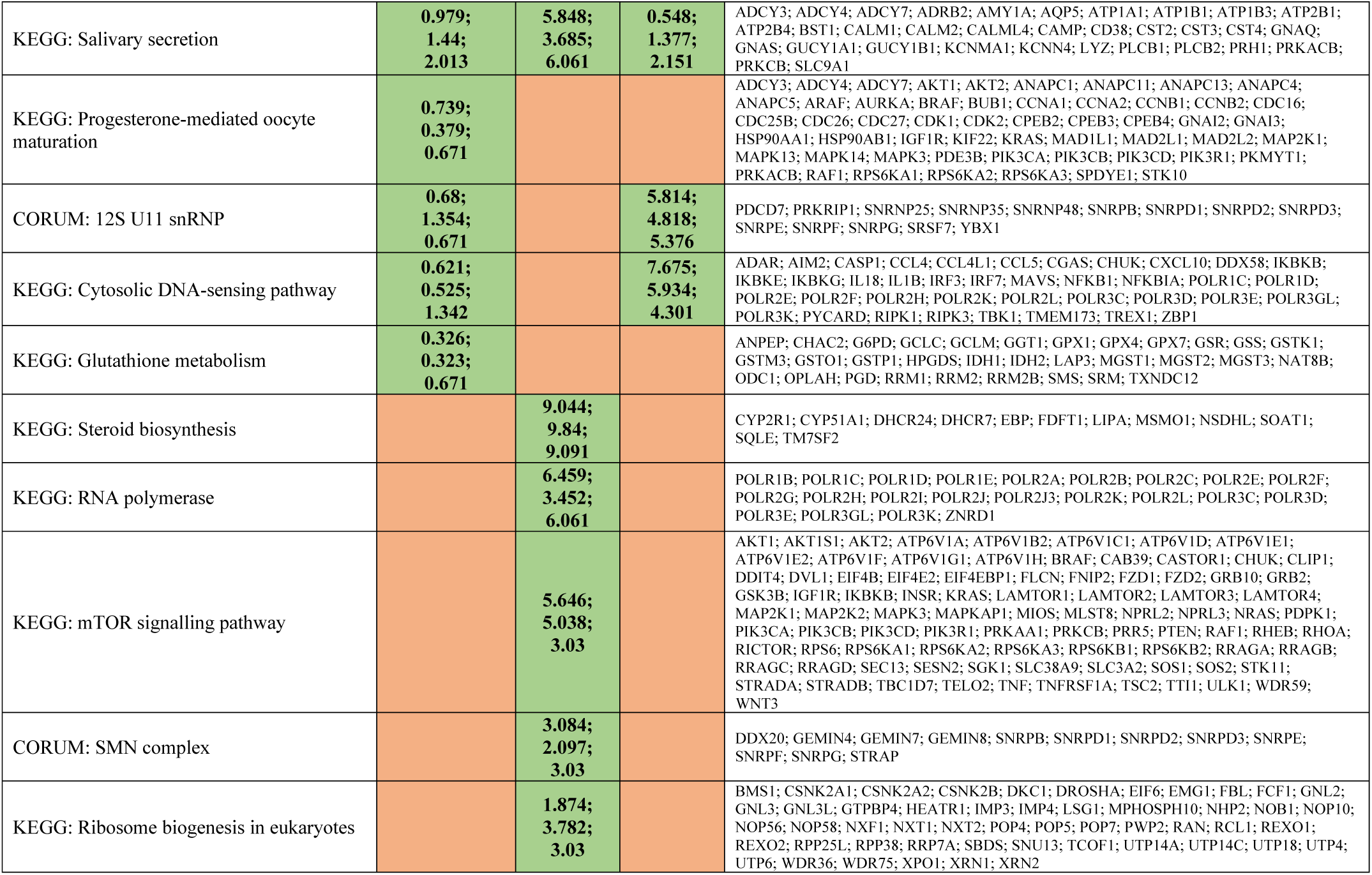
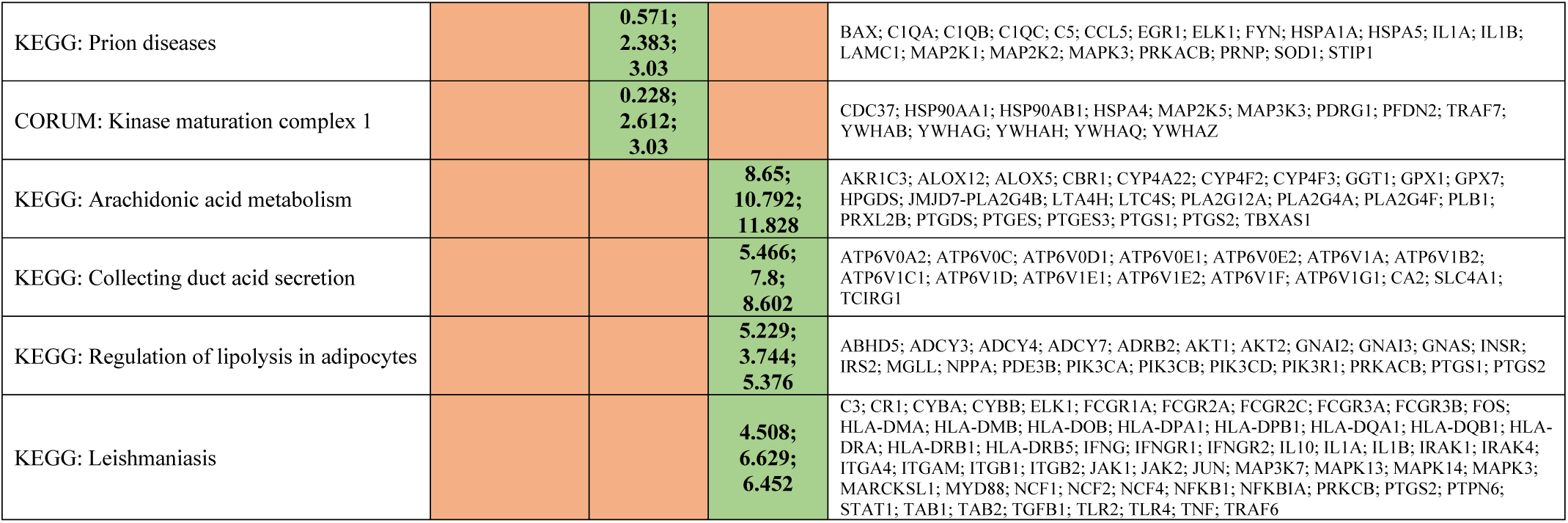
Relative contribution (gain %, cover %, frequency %) of ensemble gene noise features to the models predicting mortality/survivorship of CAP/sepsis patients

